# Skyline fossilized birth-death model is robust to violations of sampling assumptions in total-evidence dating

**DOI:** 10.1101/2023.07.23.550250

**Authors:** Chi Zhang, Fredrik Ronquist, Tanja Stadler

## Abstract

Several total-evidence dating studies under the fossilized birth-death (FBD) model have produced very old age estimates, which are not supported by the fossil record. This phenomenon has been termed “deep root attraction (DRA)”. For two specific data sets, involving divergence time estimation for the early radiations of ants, bees and wasps (Hymenoptera) and of placental mammals (Eutheria), it has been shown that the DRA effect can be greatly reduced by accommodating the fact that extant species in these trees have been sampled to maximize diversity, so called diversified sampling. Unfortunately, current methods to accommodate diversified sampling only consider the extreme case where it is possible to identify a cut-off time such that all splits occurring before this time are represented in the sampled tree but none of the younger splits. In reality, the sampling bias is rarely this extreme, and may be difficult to model properly. Similar modeling challenges apply to the sampling of the fossil record. This raises the question of whether it is possible to find dating methods that are more robust to sampling biases. Here, we show that the skyline FBD (SFBD) process, where the diversification and fossil-sampling rates can vary over time in a piecewise fashion, provides age estimates that are more robust to inadequacies in the modeling of the sampling process and less sensitive to DRA effects. In the SFBD model we consider, rates in different time intervals are either considered to be independent and identically distributed, or assumed to be autocorrelated following an Ornstein-Uhlenbeck (OU) process. Through simulations and reanalyses of the Hymenoptera and Eutheria data, we show that both variants of the SFBD model unify age estimates under random and diversified sampling assumptions. The SFBD model can resolve DRA by absorbing the deviations from the sampling assumptions into the inferred dynamics of the diversification process over time. Although this means that the inferred diversification dynamics must be interpreted with caution, taking sampling biases into account, we conclude that the SFBD model represents the most robust approach available currently for addressing DRA in total-evidence dating.

## 1 Introduction

Several studies have documented that total-evidence dating under the fossilized birth-death (FBD) model produced very old age estimates (O’Reilly et al., 2015; Zhang et al., 2016; Ronquist et al., 2016; Bapst et al., 2016; May et al., 2021). The phenomenon has been termed “deep root attraction (DRA)” (Ronquist et al., 2016), as it appears to pull age estimates towards an apparent root in the deep past. Probable reasons causing DRA include the inadequacy in morphological models failing to account for character correlations and the strong (but generally unrealistic) assumptions on diversification, fossil and extant taxa sampling in the FBD process (Ronquist et al., 2016).

Previous efforts to analyze the reasons for DRA have mainly focused on modifying the assumptions of the FBD model. It was noted that a significant improvement was obtained in analyses of both Hymenoptera (wasps, ants and bees) and Eutheria (placental mammals) data, if the model was modified to account for diversified sampling of extant taxa (Zhang et al., 2016; Ronquist et al., 2016). Diversified sampling results from the tendency of biologists to sample extant taxa that span as much phylogenetic diversity as possible (Höhna et al., 2011; Stadler and Smrckova, 2016). Diversified sampling may result in trees that are shaped quite differently than those expected from random sampling under the FBD process, in particular if only a small fraction of the total number of extant taxa is sampled. In turn, when assuming random sampling during inference while data was collected under diversified sampling, the estimated trees may be biased.

Another strategy, which has been found to be successful in reducing DRA, is to introduce strong (informative) priors on the diversification and sampling rates to penalize ghost lineages. This was shown to potentially eliminate DRA in the Eutheria data (Ronquist et al., 2016), however the priors forced some parameters of the FBD process to take on values in the posterior that are not biologically realistic. Node calibrations can also enforce strong age constraints when they are combined in total-evidence dating (Beck and Lee, 2014; O’Reilly and Donoghue, 2016), particularly when hard-bounded calibrations (such as uniform distributions) were used.

A potential factor that has not been thoroughly examined yet but that could contribute to DRA is variation in speciation, extinction and fossil-sampling rates over time in the FBD process. So far, most analyses using the FBD model have assumed that these rates are constant over time (but see Zhang et al. (2016) where the rates were shifted at two time points and May et al. (2021) where the rate in each epoch was assigned to one of the three mixture categories). For taxa which have undergone diversification for hundreds of million years, the speciation and extinction rates could have changed dramatically over time. Additionally, the fossils are usually patchy and non-uniformly sampled. Thus, assuming constant rates will impose an unrealistic prior belief on the diversification and sampling processes, and this may influence the divergence time estimates. The current paper explores to what extent variation in the FBD process rates could resolve DRA.

Specifically, we utilize the skyline fossilized birth-death (SFBD) model (Gavryushkina et al., 2014; Zhang et al., 2016), which assumes the speciation, extinction and fossil-sampling rates change through time in a piecewise constant manner. This model thus relaxes the constant rates assumption in the FBD process. Classically, in the SFBD, the rates in different time intervals are assumed to be independent and identically distributed (i.i.d.). We further developed an alternative approach assuming that rates in adjacent time intervals are similar by adding a smoothing prior to the piecewise-constant rates from the skyline model. The smoothing is done by assuming that each rate changes according to an Ornstein-Uhlenbeck (OU) process (dos Reis et al., 2012). We finally implemented the diversified sampling strategy in the SFBD model (Zhang et al., 2016) to the BDSKY package for BEAST 2 (Bouckaert et al., 2014, 2019) to extend its existing skyline functionalities. We use simulations to verify the implementation and to investigate the impact of sampling assumptions. By re-analyzing the Hymenoptera (Ronquist et al., 2012a; Zhang et al., 2016) and Eutheria data (Ronquist et al., 2016), we show that the SFBD priors resolve DRA. They also lead to robust age estimates under diversified and random sampling assumptions through their capability of inferring different process rates across time intervals, particularly the most recent intervals.

## 2 Methods

### 2.1 Skyline fossilized birth-death process

The SFBD process is conditioned on the origin time *t_or_* or the time of most recent common ancestor (MRCA) *t_mrca_*, has speciation rate *λ_i_*, extinction rate *µ_i_* and fossil-sampling rate *ψ_i_* within time interval *i* (*i* = 1, 2*, …, l*) (Gavryushkina et al., 2014; Zhang et al., 2016), with time going forward (root to tips) and *l* being the total number of time intervals. The model and implementation allow the individual rates (*λ_i_*, *µ_i_*, *ψ_i_*) to shift at different times (see Software Availability), but for convenience, we consistently specify the rates shifting at the same time in this study. Under random sampling, extant taxa are sampled uniformly at random with probability *ρ*. Under diversified sampling, there is a cutoff time, *t_c_* (i.e., *x_cut_* in Zhang et al., 2016, Fig. 1), after which only one representative extant taxon per clade is sampled. For mathematical convenience, the model further assumes no fossil sampling (*ψ_l_* = 0) and no rate shifting between *t_c_* and the present (Zhang et al., 2016). The number of sampled extant taxa (*n*; i.e. the number of extant tips in the sampled tree) and the total number of extant taxa (*N*) is an outcome of the diversified sampling process. In what follows, we refer to *ρ* = *n/N* as the sampling proportion under diversified sampling. Note that under random sampling, *ρ* is a model parameter; while under diversified sampling, *ρ* is an outcome (provided by the data), while *t_c_* is the model parameter.

**Figure 1:**
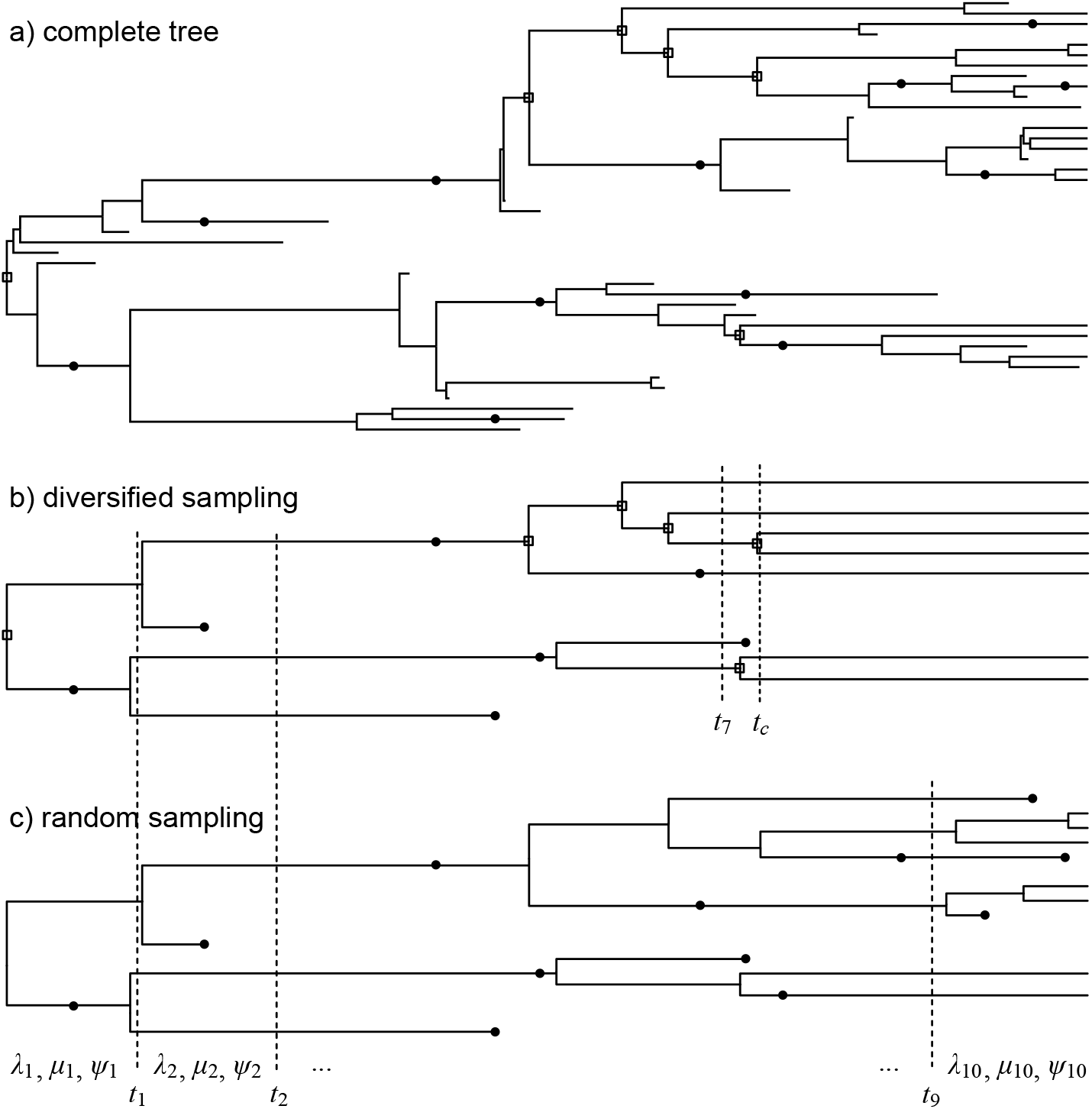
a) A tree generated from the birth-death process, with *N* = 14 extant taxa. Sampled fossils are marked as solid dots on the branches. b) For diversified sampling with proportion *ρ* = 0.5, *n* = 7 extant taxa leading to the oldest six divergences (marked as squares) in the reconstructed tree of extant taxa were selected, and no fossil was sampled more recent than the cutoff time *t_c_*. c) For random sampling, each extant taxa in the complete tree has probability *ρ* = 0.5 to be sampled. In the inference phase of the simulation study, time was divided into 10 equal-length intervals (*t*_1_ = 90, *t*_2_ = 80, . . ., *t*_9_ = 10), except for diversified sampling where the three most recent intervals were merged into one as no rate shift is assumed younger than *t_c_*.

The SFBD process provides the probability density of a tree with extant and fossil samples given the model parameters. This probability density is given in Zhang et al. (2016, Eqn 1 for random sampling and Eqn 4 for diversified sampling). For diversified extant sampling, the proportion *ρ* (or the total number of extant taxa, *N*) has to be additionally supplied as data when calculating the probability density of the sampled tree (Zhang et al., 2016, Eqn 7). Throughout this work, we provide *ρ* and evaluate *N* through *ρ* = *n/N* . In our implementations of diversified sampling, the parameter *t_c_* is placed slightly after the last split, fossil, or rate-shifting time, whichever is the youngest, to meet the requirement of the model, and is updated during Markov chain Monte Carlo (MCMC) computation. Under both sampling schemes, *ρ* is typically fixed based on biological knowledge, while we estimate the remaining parameters (i.e., *λ_i_*, *µ_i_*, and *ψ_i_*).

In Bayesian total-evidence dating under the SFBD process, we have to specify the priors for the rates *λ_i_*, *µ_i_* and *ψ_i_*. When the rates are assumed to be i.i.d. across time intervals, the same prior probability distribution is typically assigned for each rate (e.g., an exponential distribution). As the number of time intervals increases, the available data (i.e., bifurcations and fossils in the tree) decreases in each interval, making it difficult to infer independent rates reliably. To address this, we can take advantage of the fact that rates (*λ_i_*, *µ_i_*or *ψ_i_*) tend to be autocorrelated between intervals, especially when intervals are short. Such correlation have been modeled by the Brownian motion (Silvestro et al., 2019) and more regularizing Horseshoe Markov random field (Magee et al., 2020). We alternatively model the rates using the OU process (Uhlenbeck and Ornstein, 1930; du Plessis, 2016).

### 2.2 OU process prior

The OU process is a univariate Markov process. It reverts to the long-term mean *θ* with strength *ν* after any perturbation. Upon reaching the mean, it fluctuates around it with variance *σ*^2^. Due to its memoryless property, the likelihood of a trajectory of points, *x*_0_*, x*_1_*, …, x_l_*, is the product of normal distributions:

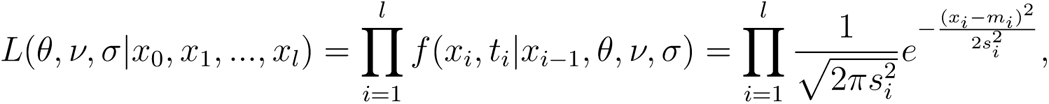

where *m _i_* = *θ* +(*x_i_*_−1_ − *θ*)*e*^−*νti*^ is the mean, 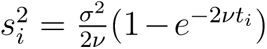 is the variance, and *t_i_* is the time duration of interval *i*. Replacing *x_i_* with *λ_i_*, *µ_i_*or *ψ_i_*, we obtain the prior distribution for each parameter.

The implementation of the OU process prior was by reusing the code written by du Plessis (2016). It contains three hyperparameters for each rate, *θ*, *ν* and *σ*, that need to be assigned hyperpriors. The process is less intuitive than the i.i.d. case, thus we provide an R script for simulating the OU trajectories to help choosing the hyperpriors (Supplementary Material).

### 2.3 Simulations

To verify the implementation of diversified extant-sampling and the related skyline functionalities in the BDSKY package, we performed a series of computer simulations. The parameters are chosen to generate reasonably large trees and data sets that are computationally tractable, and also in the same magnitude as the empirical data analyzed later.

#### Generating birth-death trees

Using TreeSim in R (Stadler, 2011), we generated 200 rooted trees from the birth-death process with constant birth rate (*λ*) of 0.08 and death rate (*µ*) of 0.05, conditioned on the root age of 100 Ma. These trees have the number of extant tips (*N*) ranging from 8 to 362 (median = 90) and extinct tips ranging from 12 to 546 (median = 136)

#### Sampling extant taxa

Given each birth-death tree, we sampled extant taxa using either random or diversified sampling. For random sampling, each extant tip has equal probability (*ρ*) of 0.5 to be sampled. For diversified sampling, we set *t_c_* to the round(*N/*2 − 1)’th oldest divergence in the extant-taxa tree (Fig. 1), meaning the sampled tree has half as many extant tips compared to the full tree (i.e., sampling proportion *ρ* = 0.5).

#### Sampling fossils

Fossils were sampled with constant rate (*ψ*) of 0.02 along the branches of the birth-death tree, from the root to the present under random sampling, or from the root to time *t_c_* under diversified sampling (*ψ* = 0 after *t_c_*). Unsampled lineages were pruned from the complete tree to obtain the sampled tree (Fig. 1). The resulting trees have 7 to 222 fossils (median = 55) under random sampling, and 5 to 147 fossils (median = 38) under diversified sampling.

#### Generating morphological characters

Given each sampled tree, we simulated discrete morphological matrices under the Markov k-states variable (Mkv) model (Lewis, 2001), that is, keeping only the variable characters in the matrices. Each data matrix contains 200 characters from binary to up to five states with proportions of 0.4, 0.3, 0.2 and 0.1, respectively.

We used either a constant evolutionary rate of 0.02 per character per myr (strict clock), or a variable evolutionary rate modeled by the uncorrelated lognormal distributions (UCLD) relaxed clock (Drummond et al., 2006). For the latter, the mean rate was 0.02 per character per myr, while the relative rate multiplier for each branch was drawn from a lognormal distribution with mean 1.0 and variance 1.0 (both in real space) (cf. Zhang, 2021, Fig. 1c).

Extant-species and fossil sampling and morphological-character simulation were performed using a self-written C program. The source code and simulation scripts are provided in Supplementary Material.

#### Inference from fixed tree

In the first round of simulations, we fixed the tree (both topology and node ages) in the inference to the simulated one to infer the SFBD rates (*λ_i_*, *µ_i_* and *ψ_i_*). Time was divided into 10 equal-length intervals (i.e., *l* = 10 and the rate-shifting times are 90, 80, . . ., 10 Ma), except for inferences under diversified sampling. To avoid the last shifting time to be younger than the cutoff time *t_c_*, the three most recent intervals were merged (i.e., *l* = 8 and the youngest rate-shifting time is 30 Ma). We tested cases when the sampling strategy and proportion (*ρ*) were set to the truth, but also cases when these sampling assumptions were mismatched between simulation and inference. The details of all 16 cases are listed in Table 1.

**Table 1:** Parameter values used in the simulations, and SFBD model and prior settings in the subsequent inferences. Inference 1, 3, 9 and 11 present the best match to the simulation setting. Abbreviations: rnd: random sampling; div: diversified sampling; itv: time intervals (*i* = 1*, . . ., l*).

|  |  |
| --- | --- |
| Simulation: | $\lambda = 0.08, \mu = 0.05, \psi = 0.02, \rho = 0.5, \text{rnd}$ |
| Inference: 1 | i.i.d. 10 itv, $\lambda_i, \mu_i, \psi_i \sim \text{Exp}(10), \rho = 0.5, \text{rnd}$ |
| 2 | i.i.d. 10 itv, $\lambda_i, \mu_i \sim \text{Exp}(1), \psi_i \sim \text{Exp}(10), \rho = 0.5, \text{rnd}$ |
| 3 | OU, 10 itv, $\lambda_0, \mu_0, \psi_0 \sim \text{Exp}(10), \rho = 0.5, \text{rnd}$ |
| 4 | OU, 10 itv, $\lambda_0, \mu_0 \sim \text{Exp}(1), \psi_0 \sim \text{Exp}(10), \rho = 0.5, \text{rnd}$ |
| 5 | i.i.d. 10 itv, $\lambda_i, \mu_i, \psi_i \sim \text{Exp}(10), \rho = 0.1, \text{rnd}$ |
| 6 | i.i.d. 10 itv, $\lambda_i, \mu_i, \psi_i \sim \text{Exp}(10), \rho = 1.0 \text{ complete}$ |
| 7 | i.i.d. 10 itv, $\lambda_i, \mu_i, \psi_i \sim \text{Exp}(10), \rho = 0.5, \text{div}$ |
| 8 | OU, 10 itv, $\lambda_0, \mu_0, \psi_0 \sim \text{Exp}(10), \rho = 0.5, \text{div}$ |
| Simulation: | $\lambda = 0.08, \mu = 0.05, \psi = 0.02, \rho = 0.5, \text{div}$ |
| Inference: 9 | i.i.d. 8 itv, $\lambda_i, \mu_i, \psi_i \sim \text{Exp}(10), \rho = 0.5, \text{div}$ |
| 10 | i.i.d. 8 itv, $\lambda_i, \mu_i \sim \text{Exp}(1), \psi_i \sim \text{Exp}(10), \rho = 0.5, \text{div}$ |
| 11 | OU, 8 itv, $\lambda_0, \mu_0, \psi_0 \sim \text{Exp}(10), \rho = 0.5, \text{div}$ |
| 12 | OU, 8 itv, $\lambda_0, \mu_0 \sim \text{Exp}(1), \psi_0 \sim \text{Exp}(10), \rho = 0.5, \text{div}$ |
| 13 | i.i.d. 8 itv, $\lambda_i, \mu_i, \psi_i \sim \text{Exp}(10), \rho = 0.1, \text{div}$ |
| 14 | i.i.d. 10 itv, $\lambda_i, \mu_i, \psi_i \sim \text{Exp}(10), \rho = 1.0 \text{ complete}$ |
| 15 | i.i.d. 10 itv, $\lambda_i, \mu_i, \psi_i \sim \text{Exp}(10), \rho = 0.5, \text{rnd}$ |
| 16 | OU, 10 itv, $\lambda_0, \mu_0, \psi_0 \sim \text{Exp}(10), \rho = 0.5, \text{rnd}$ |

We used exponential priors for the i.i.d. rates. For the OU process rates, we used the same exponential hyperpriors for *x*_0_ and *θ*, and set Gamma(4, 0.05) (mean = 0.2) for *σ* and Gamma(4, 0.5) (mean = 2) for *ν* (the gamma distribution used the shape and scale parameterization). These hyperpriors were chosen based on the simulated trajectories, which appear to have sufficient variation and do not inappropriately constrain potential changes in process rates over time (Supplementary Fig. S1).

The MCMC chain was run 5 million generations and sampled every 500 generations, with the beginning 1/4 samples discarded as burn-in. Convergence and sufficient sampling of parameters were assessed through trace plots and effective sample sizes (ESS *>* 200).

#### Inference from morphological data

In the following two rounds, we used the morphological data simulated under strict and relaxed clock models for inference. The evolutionary model is Mkv matching the simulation setting. The trees were co-estimated rather than fixed. The fossil ages were fixed to their true values and root age was assigned a broad Uniform(0, 200) prior. Under strict clock, the evolutionary rate was given an Uniform(0, 1) prior; while under the relaxed clock, the mean rate had Uniform(0, 1) prior and the standard deviation parameter had Exp(1.0) prior. The rest of the prior settings follow the previous section (Table 1).

The MCMC chain length was increased to 20 million to guarantee sufficient ESS. Topological convergence was assessed by the sliding-window and cumulative average of split frequencies using the R package RWTY (Warren et al., 2017).

### 2.4 Hymenoptera

We re-analyzed the Hymenoptera data consisting of 113 taxa (68 extant species and 45 fossils), 353 morphological characters and 6 molecular genes (Ronquist et al., 2012a; Zhang et al., 2016), under the SFBD model with i.i.d. and OU process rates. The fossil ages were updated according to O’Reilly and Donoghue (2016) (Supplementary Table S1).

We focused on the SFBD model with time divided into either *l* = 10 (shifting at 280, 250, . . ., 70, 40 Ma) or 20 (shifting at 310, 295, 280, . . ., 70, 55, 40 Ma) intervals (Table 2). For i.i.d. rates, the prior was Exp(10) (mean = 0.1) for all *λ_i_*, *µ_i_* and *ψ_i_*, or was Exp(10) for *d_i_* and Uniform(0, 1) for *r_i_* and *s_i_* (which convert to *λ_i_* = 0.2, *µ_i_*= *ψ_i_* = 0.1 if using the means of these distributions). A mean rate of 0.1 means the expected waiting time for an event (a lineage to diverge, to go extinct, or to sample a fossil) is 1/0.1 = 10 Myr. We only used the *λ*, *µ*, *ψ* parameterization under the OU prior. Equivalent to the simulations, we used Exp(10) for *x*_0_ and *θ*, Gamma(4, 0.05) for *σ*, and Gamma(4, 0.5) for *ν* as hyperpriors for {*λ_i_*}, {*µ_i_*} and {*ψ_i_*}. We also further relaxed the variance under 20 intervals, using a Gamma(4, 0.2) distribution (mean = 0.8) for *σ*.

**Table 2:** SFBD model and prior settings in the Hymenoptera and Eutheria analyses. Abbreviations: div: diversified sampling; rnd: random sampling; itv: time intervals (*i* = 1*, . . ., l*).

|  |  |
| --- | --- |
| Hymenoptera: |  |
| 1 | Constant, $\lambda, \mu, \psi \sim \text{Exp}(10)$, $\rho = 0.001$, div & rnd |
| 2 | Constant, $d \sim \text{Exp}(10)$, $r, s \sim \text{U}(0, 1)$, $\rho = 0.001$, div & rnd |
| 3 | i.i.d. 10 itv, $d_i \sim \text{Exp}(10)$, $r_i, s_i \sim \text{U}(0, 1)$, $\rho = 0.001$, div & rnd |
| 4 | i.i.d. 10 itv, $\lambda_i, \mu_i, \psi_i \sim \text{Exp}(10)$, $\rho = 0.001$, div & rnd |
| 4b | i.i.d. 10 itv, $\lambda_i, \mu_i, \psi_i \sim \text{Exp}(10)$, $\rho = 0.1$, div |
| 5 | i.i.d. 20 itv, $\lambda_i, \mu_i, \psi_i \sim \text{Exp}(10)$, $\rho = 0.001$, div & rnd |
| 6 | OU, 10 itv, $\lambda_0, \mu_0, \psi_0 \sim \text{Exp}(10)$, $\rho = 0.001$, div & rnd |
| 6b | OU, 10 itv, $\lambda_0, \mu_0, \psi_0 \sim \text{Exp}(10)$, $\rho = 0.1$, div |
| 7 | OU, 20 itv, $\lambda_0, \mu_0, \psi_0 \sim \text{Exp}(10)$, $\rho = 0.001$, div & rnd |
| Eutheria: |  |
| 1 | Constant, $\lambda, \mu, \psi \sim \text{Exp}(10)$, $\rho = 0.01$, div & rnd |
| 2 | Constant, $d \sim \text{Exp}(10)$, $r, s \sim \text{U}(0, 1)$, $\rho = 0.01$, div & rnd |
| 3 | i.i.d. 10 itv, $d_i \sim \text{Exp}(10)$, $r_i, s_i \sim \text{U}(0, 1)$, $\rho = 0.01$, div & rnd |
| 4 | i.i.d. 10 itv, $\lambda_i, \mu_i, \psi_i \sim \text{Exp}(10)$, $\rho = 0.01$, div & rnd |
| 4b | i.i.d. 10 itv, $\lambda_i, \mu_i, \psi_i \sim \text{Exp}(10)$, $\rho = 0.2$, div |
| 5 | i.i.d. 20 itv, $\lambda_i, \mu_i, \psi_i \sim \text{Exp}(10)$, $\rho = 0.01$, div & rnd |
| 6 | OU, 10 itv, $\lambda_0, \mu_0, \psi_0 \sim \text{Exp}(10)$, $\rho = 0.01$, div & rnd |
| 6b | OU, 10 itv, $\lambda_0, \mu_0, \psi_0 \sim \text{Exp}(10)$, $\rho = 0.2$, div |
| 7 | OU, 20 itv, $\lambda_0, \mu_0, \psi_0 \sim \text{Exp}(10)$, $\rho = 0.01$, div & rnd |

The root age (*t_mrca_*) was assigned an exponential prior with mean 81 and offset 315 Ma (referring to *Katerinka* (oldest Neoptera) and *Rhyniognatha* (oldest insect), same as in Ronquist et al. (2012a); Zhang et al. (2016)). The Holometabola clade was constrained to be monophyletic to root the tree properly (Ronquist et al., 2012a; Zhang et al., 2016), but different from the previous studies, we did not add a calibration distribution for the Holometabola clade. The extant sampling proportion (*ρ*) was fixed to 0.001 for both random and diversified sampling (meaning 60 out of approximately 60,000 extant hymenopteran species were sampled). For the UCLD clock model, we used a weakly informative Lognormal(−7, 0.5) prior (parameterized on the log scale such that the median on the natural scale is *e*^−7^) for the mean rate and an Exp(1.0) prior for the standard deviation, based on the previous studies (Ronquist et al., 2012a; Zhang et al., 2016).

Each of two independent MCMC chains was run 200 million generations and sampled every 5000 generations. The first 30% samples were discarded as burn-in. Convergence and sufficient sampling of parameters were assessed through trace plots and ESS values (*>* 200), while topological convergence was assessed by the sliding-window (within a run) and average standard deviation of split frequencies (ASDSF *<* 0.02 among runs) using RWTY in R (Warren et al., 2017).

### 2.5 Eutheria

We also re-analyzed the Eutheria data assembled in Ronquist et al. (2016), which contains 74 taxa (33 fossils and 41 extant species), 3998 morphological characters (keeping only the variable ones out of 4541 in total), and 36860 nucleotide sites (27 nuclear genes divided into 4 partitions).

Most of the settings follow the Hymenoptera analyses unless mentioned below (Table 2). The rate-shifting times are 110, 100, . . ., 40, 30 Ma for *l* = 10, and are 120, 110, 105, . . ., 35, 30, 25 Ma for *l* = 20. The origin time (*t_or_*) was assigned an exponential prior with mean 45 and offset 100 Ma (referring to *Eomaia scansoria*, Supplementary Table S2. The mean, 145 Ma, of this offset-exponential distribution is the boundary of Jurassic and Cretaceous.). The extant sampling proportion (*ρ*) was fixed to 0.01 for both random and diversified sampling, based on the number of living placental mammal species. We used a weakly informative Lognormal(−6, 0.5) prior (with median *e*^−6^) for the mean rate in the UCLD clock model following the previous study (Ronquist et al., 2016).

Each chain was run 400 million generations and sampled every 5000 generations, and the first 50% samples were discarded as burn-in. The Boreoeutheria (Laurasiatheria + Euarchontoglires) clade was constrained to be monophyletic to correct the rooting (Ronquist et al., 2016). To overcome bimodal posterior of the node age and to achieve good convergence and mixing (particularly under random sampling), the MRCA of Boreoeutheria was assigned a Normal(80, 10^2^) distribution based on information from previous analyses (Ronquist et al., 2016). This distribution is relatively broad as the 95% density ranges from 60 to 100, which is about two times wider than the posterior CI from Ronquist et al. (2016) under various prior settings. (The runs under diversified sampling converged well without this calibration, but this calibration was also used to be consistent.)

Complete prior and MCMC settings can be found in the XML files in Supplementary Material.

## 3 Results

### 3.1 Simulations

#### Inference from fixed tree

We first inferred the SFBD rates when the input trees (both topologies and node ages) were fixed to the true trees in the simulations. Under these conditions, the only estimated parameters are *λ_i_*, *µ_i_*, and *ψ_i_* (*i* = 1, . . ., *l*). We summarized the posterior medians (Figs 2– 5), and the relative widths of the 95% highest posterior density credibility interval (HPD CI, the width divided by the true value, Supplementary Figs S2–S5), of each parameter under each of the 16 conditions in Table 1.

**Figure 2:**
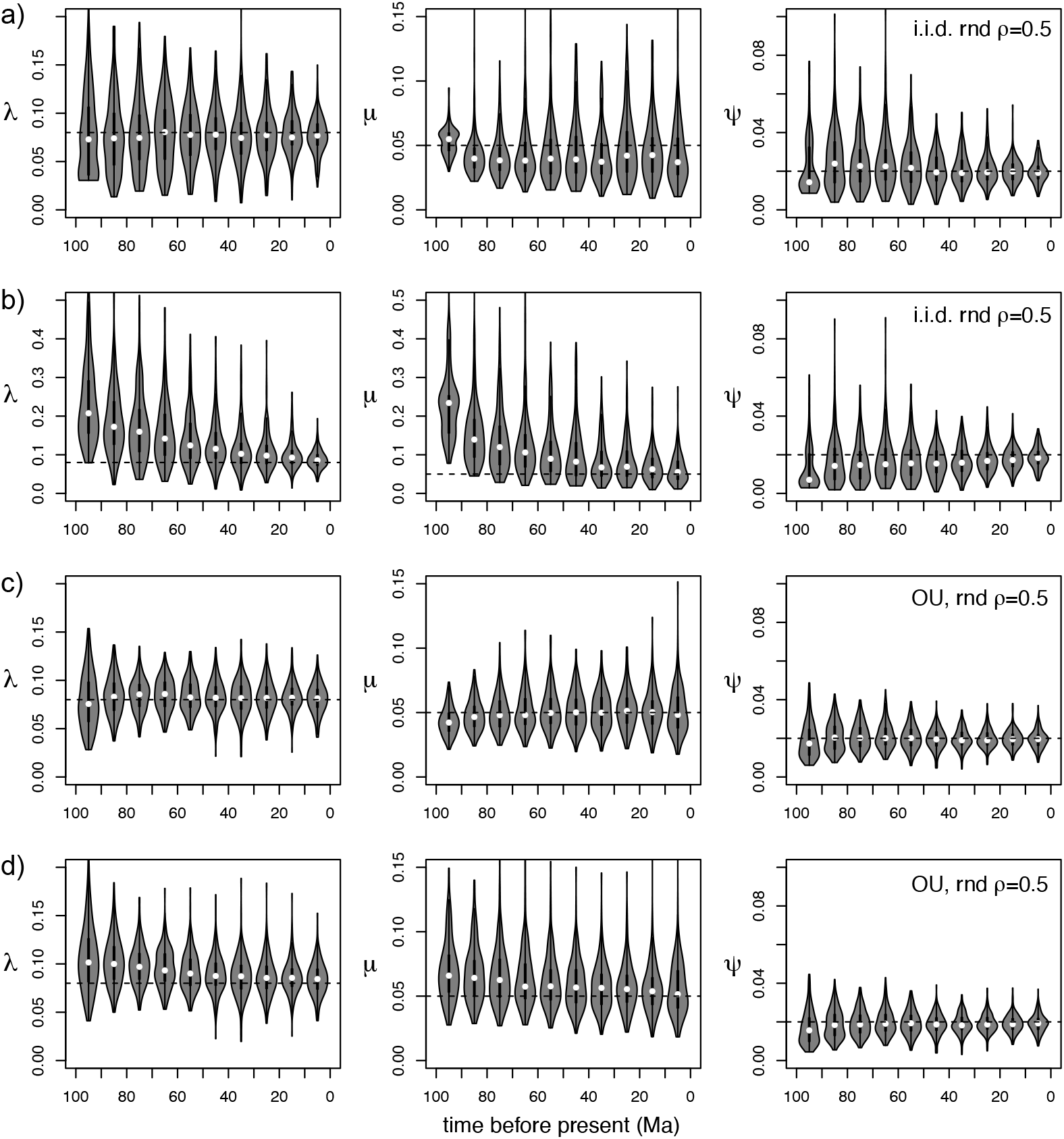
Posterior medians for the SFBD rates under various conditions in Table 1: a) Inference 1; b) Inference 2; c) Inference 3; d) Inference 4. Each violin plot summarizes the values across 200 replicates (i.e., for 200 trees). Input trees were fixed to the true trees simulated under random sampling (rnd), with sampling assumptions matched between simulation and inference. The doted horizontal line represents the true value of the parameter. See Supplementary Figure S2 for relative CI widths.

**Figure 5:**
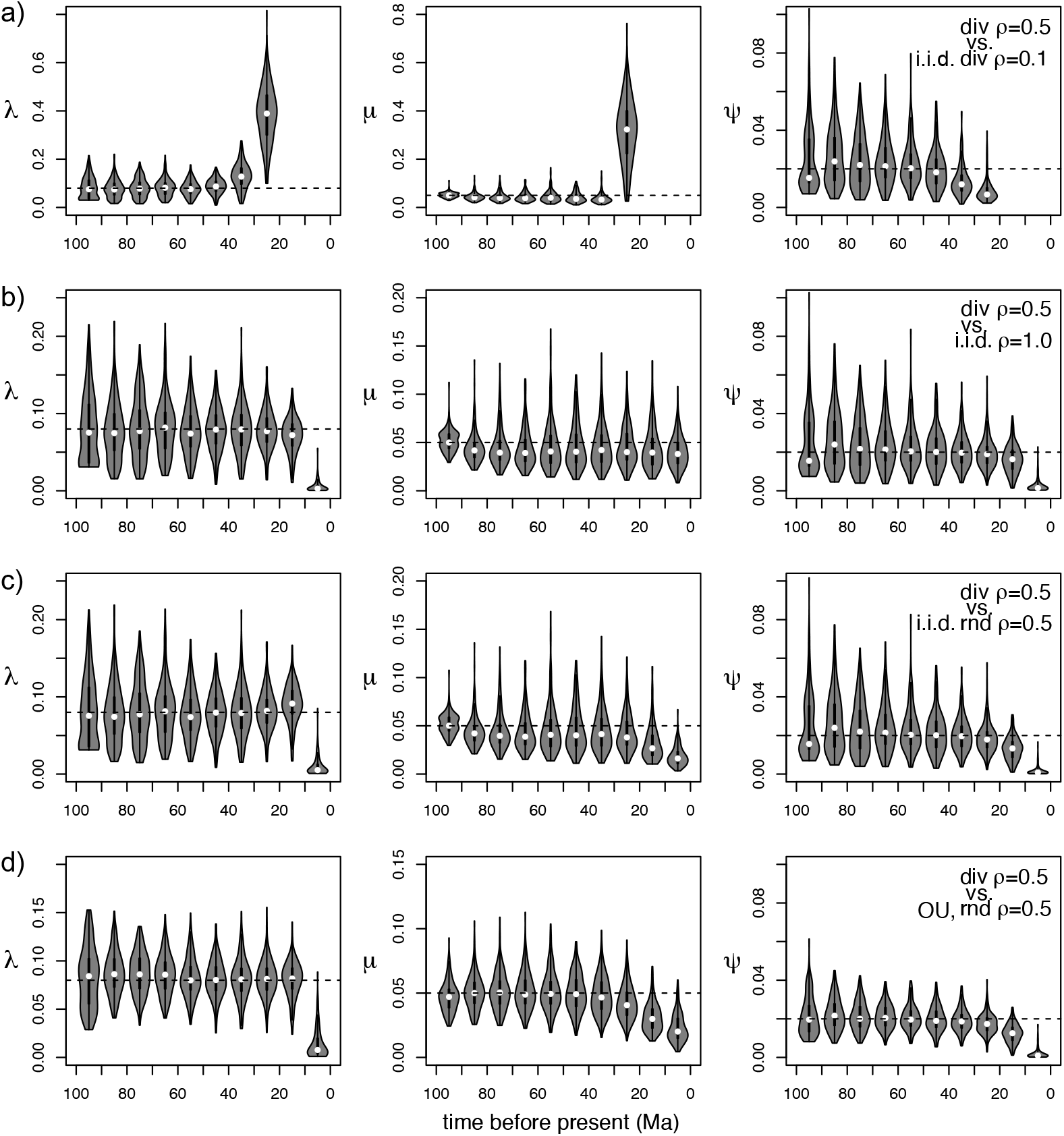
Posterior medians for the SFBD rates under various conditions in Table 1: a) Inference 13; b) Inference 14; c) Inference 15; d) Inference 1. 16. Each violin plot summarizes the values across 200 replicates (i.e., for 200 trees). Input trees were fixed to the true trees under diversified sampling (div), but sampling assumptions were mismatched between simulation and inference. The doted horizontal line represents the true value of the parameter. See Supplementary Figure S5 for relative CI widths.

When the sampling assumptions were matched between simulation and inference, and the prior mean was close to the true value of the parameter, the posterior medians are also close to the true value (Fig. 2ac, Fig. 4ac, and Supplementary Figs S2 & S4), although the CIs are relatively wider at older time intervals then at younger ones (Supplementary Figs S2 & S4).

When the prior mean was more than 10 times of the true value, the posterior medians are much larger than the true value for the oldest intervals (Fig. 2bd and Fig. 4bd, for *λ* and *µ*) with much wider CIs (Supplementary Figs S2 & S4), due to the fact that the number of lineages is small near the root of the tree. Comparing with the i.i.d. rate prior, the OU prior helped with reducing the overestimation of birth and death rates through its smoothing effect (Figs 2 and 4, Supplementary Figs S2 & S4).

**Figure 4:**
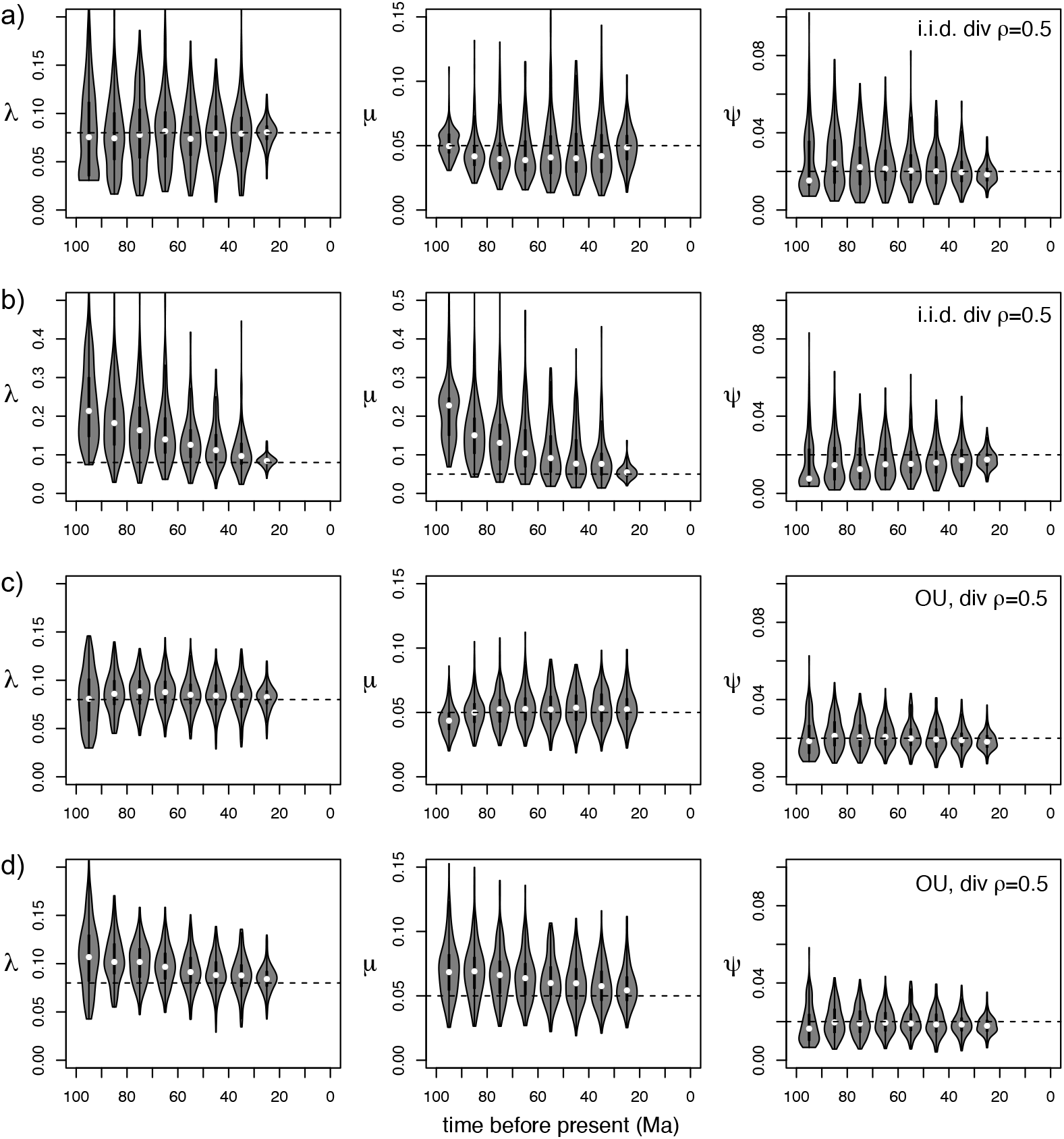
Posterior medians for the SFBD rates under various conditions in Table 1: a) Inference 9; b) Inference 10; c) Inference 11; d) Inference 12. Each violin plot summarizes the values across 200 replicates (i.e., for 200 trees). Input trees were fixed to the true trees under diversified sampling (div), with sampling assumptions matched between simulation and inference. The doted horizontal line represents the true value of the parameter. Note the last interval ranges from 30 to 0. See Supplementary Figure S4 for relative CI widths.

When a smaller sampling proportion (0.1), larger sampling proportion (1.0, complete sampling), or different sampling strategy (random vs. diversified) was used in the inference, the posterior rate estimates for the most recent intervals are greatly affected, but the estimates at the oldest five or more intervals are barely affected (Figs 3 and 5, Supplementary Figs S3 & S5). Particularly, the speciation and extinction rates in the youngest interval are largely elevated under diversified sampling when the trees were simulated under random sampling (Fig. 3cd), or the diversified sampling was matched but under a smaller proportion (Fig. 5a).

**Figure 3:**
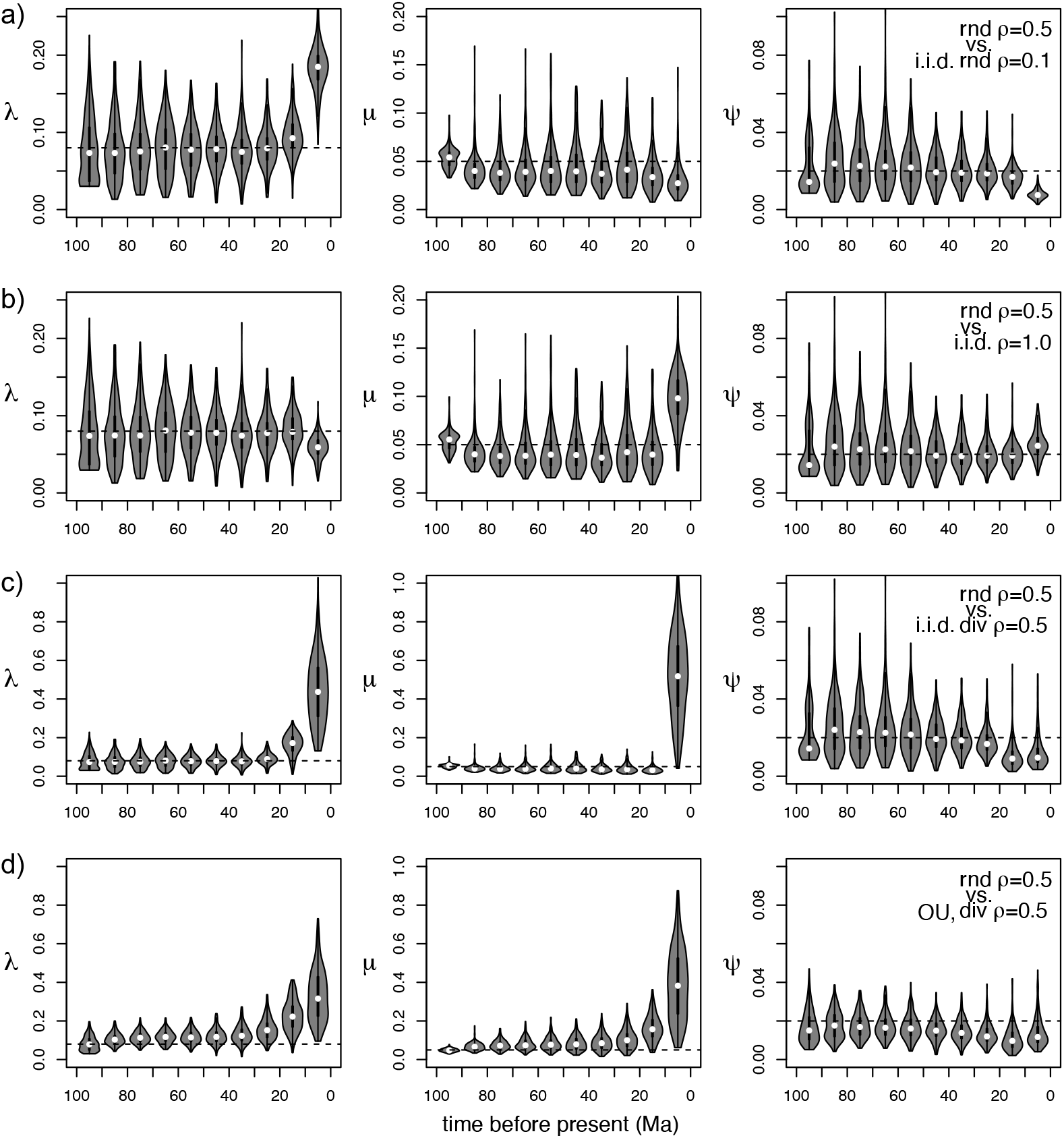
Posterior medians for the SFBD rates under various conditions in Table 1: a) Inference 5; b) Inference 6; c) Inference 7; d) Inference 8. Each violin plot summarizes the values across 200 replicates (i.e., for 200 trees). Input trees were fixed to the true trees under random sampling (rnd), but sampling assumptions were mismatched between simulation and inference. The doted horizontal line represents the true value of the parameter. See Supplementary Figure S3 for relative CI widths.

#### Inference from morphological data

We also inferred the parameters (including the trees) directly from the simulated morphological characters. The Mkv model was consistently used and no missing character was involved. We summarized the posterior medians and relative CI widths of the tree height and (mean) clock rate, as well as the normalized Robinson-Foulds (RF) distances between each MCC tree and the true tree, under each of the 16 conditions (Table 1). Figure 6 shows the results when the data were simulated under the strict clock, while Figure 7 is under the relaxed clock.

**Figure 6:**
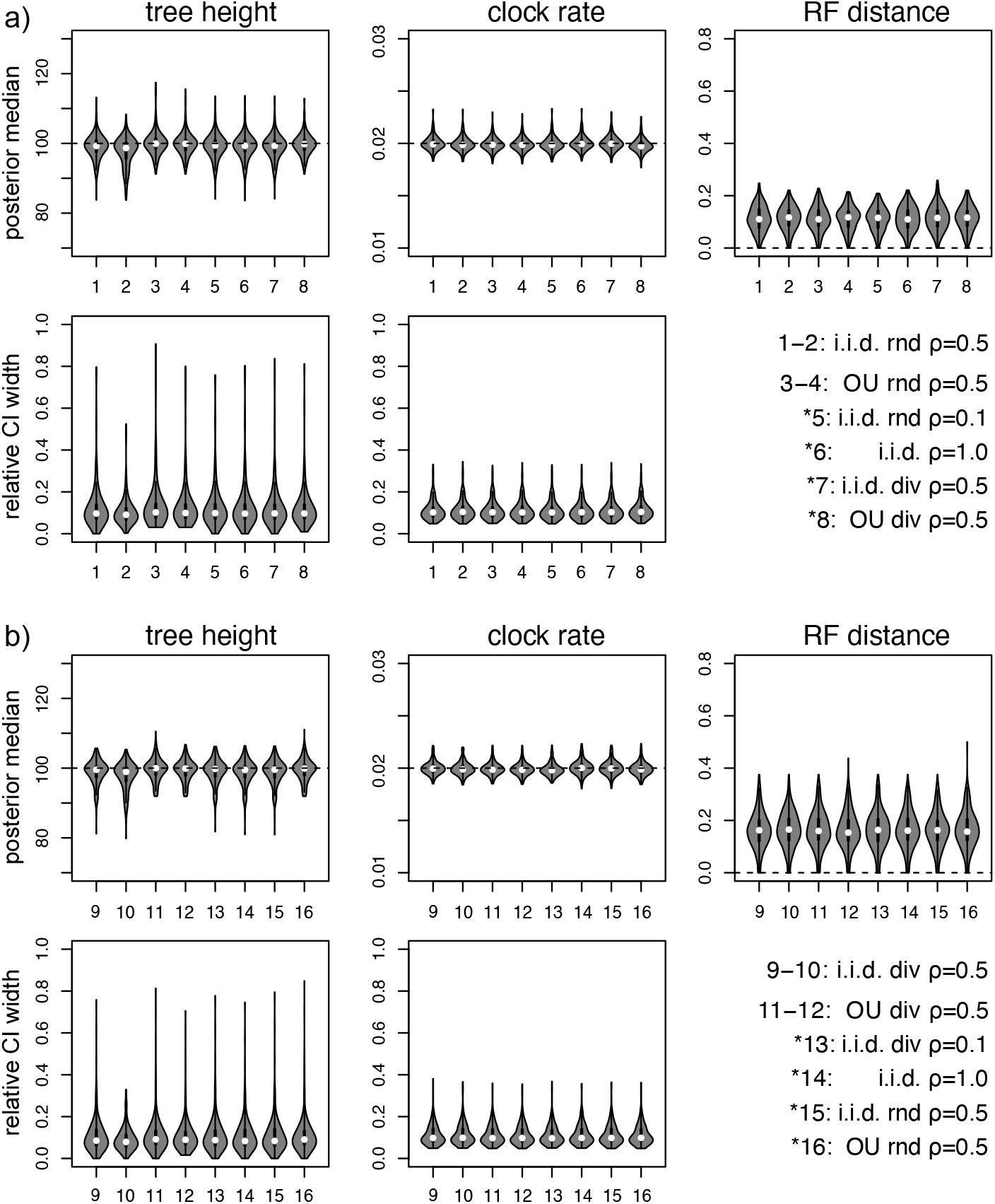
Posterior medians for the tree height and clock rate, and normalized RF distances, followed by the relative CI widths for the tree height and clock rate, under each of the 16 conditions in Table 1. Each violin plot summarizes the values across 200 replicates. Morphological matrices were simulated under strict clock given trees sampled under a) random sampling and b) diversified sampling. Inferences marked with a star have mismatched extant sampling assumptions. The doted horizontal line represents the true value of the parameter.

**Figure 7:**
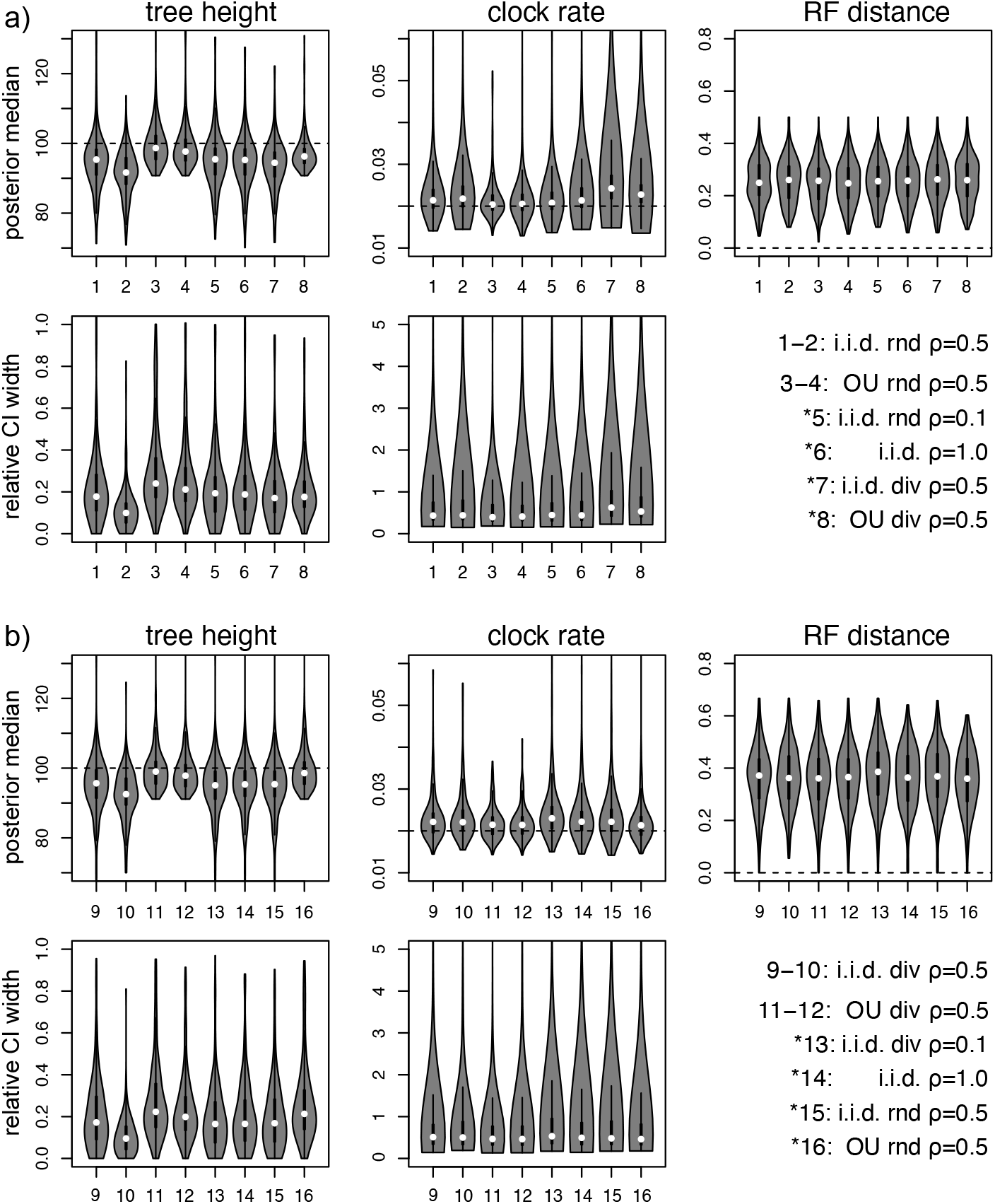
Posterior medians for the tree height and mean clock rate, and normalized RF distances, followed by the relative CI widths for the tree height and mean clock rate, under each of the 16 conditions in Table 1. Each violin plot summarizes the values across 200 replicates. Morphological matrices were simulated under UCLD relaxed clock given trees sampled under a) random sampling and b) diversified sampling. Inferences marked with a star have mismatched extant sampling assumptions. The doted horizontal line represents the true value of the parameter.

The posterior estimates of the tree height and clock rate are quite robust to various SFBD assumptions, showing good accuracy and precision under these conditions (Fig. 6). The morphological data appear very informative to produce a reliable estimate of the tree, as a result, the SFBD rates show similar patterns to when the tree was fixed (Supplementary Figs S6–S13). The parameter estimates become less reliable under the relaxed clock, showing more variable medians among different conditions and much wider CIs (Fig. 7, see also Supplementary Figs S14–S21 for the SFBD rates). The most accurate results are produced under the OU process priors, while the least ones are under the i.i.d. Exp(1) priors for the speciation and extinction rates as most uncertainties were involved.

In summary, the simulation results demonstrate the ability of the SFBD implementation recovering the true parameter values under an appropriate sampling assumption, and the robustness of root age estimates to violations of certain sampling assumptions.

For both the Hymenoptera and Eutheria data, the i.i.d. and OU priors for the SFBD rates resolve DRA and unify the age estimates under random and diversified sampling assumptions. The age estimates are quite consistent across these prior settings. In contrast, under the constant FBD rates, the age estimates were estimated in previous studies to be much older under random sampling then those under diversified sampling (Zhang et al., 2016; Ronquist et al., 2016), with differences in some details. We discuss each data set in more detail in the following sections.

### 3.2 Hymenoptera

As a cross validation of the implementations between BEAST 2 and MrBayes 3.2 (Ronquist et al., 2012b), we first checked the consistence of parameter estimates under both strict and UCLD relaxed clocks, while trying to match all possible settings in both programs. The results cross validate the implementation of the SFBD model in both programs (Supplementary Tables S3–S5).

When the FBD rates are constant, the age estimates of the Hymenoptera radiation are dramatically different under random and diversified sampling assumptions (Fig. 8, first 2 bars). In the *λ, µ, ψ* parameterization, the five (median) representative ages are more than doubled under the random-sampling assumption compared to the diversified-sampling assumption. Linked to this, the base clock rate under the random-sampling assumption is inferred to be only 1/3 of the rate under the diversified-sampling assumption. In the *d, r, s* parameterization, DRA under random sampling is less prominent but still obvious. Since the oldest hymenopteran fossils, *Triassoxyela* and *Asioxyela*, are dated to the Carnian and Ladinian in the Triassic (228–242 Ma), a crown age of Hymenoptera in the Permian (252–299 Ma) appears plausible, while a crown age of Hymenoptera in the Carboniferous (299–359 Ma) or older would leave a big gap (thus long ghost lineages) to these fossils.

**Figure 8:**
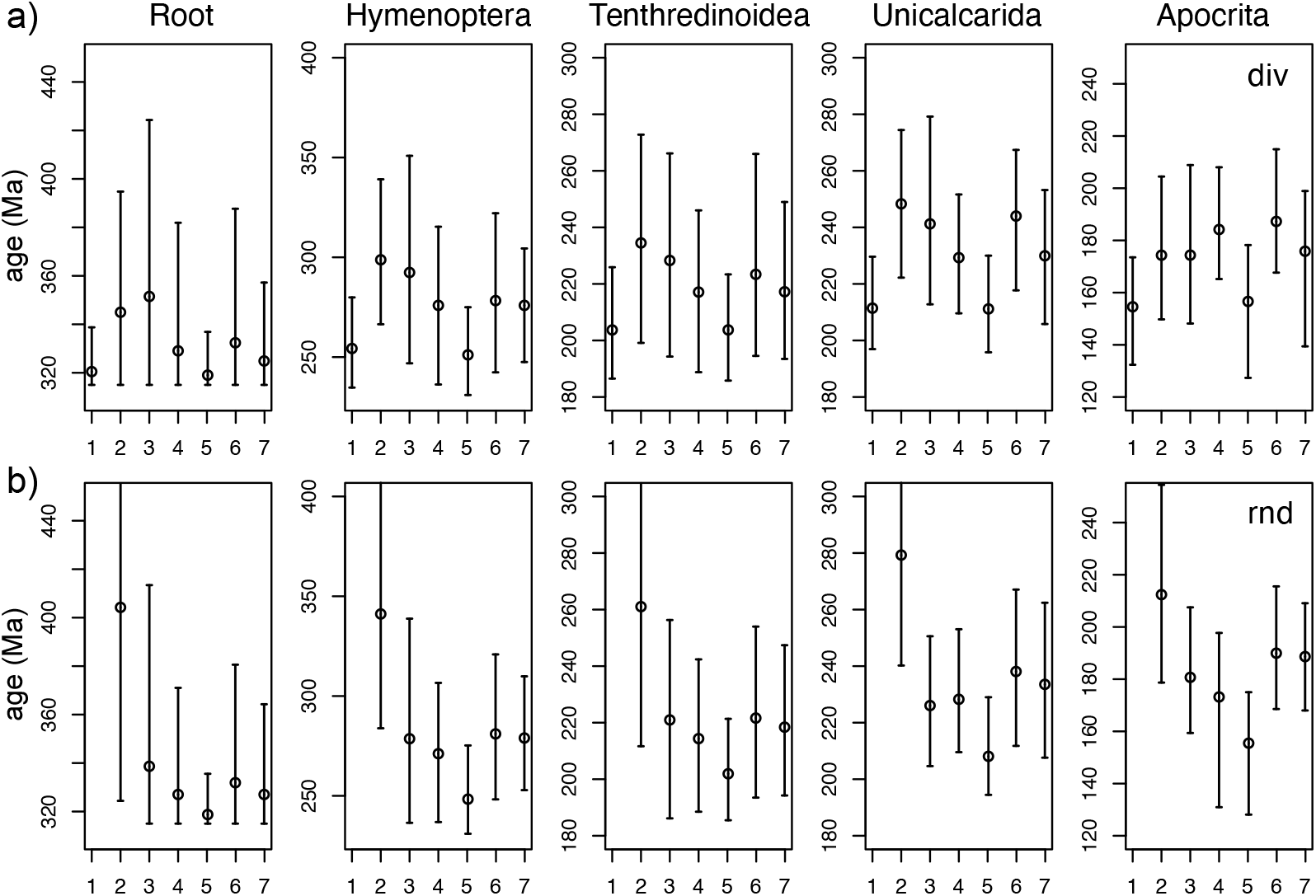
Posterior estimates of five representative node ages (Ma, median and 95% HPD CI) in the Hymenoptera analyses, using the SFBD model settings shown in Table 2 under a) diversified sampling and b) random sampling. The first two cases in each panel are under constant-rate FBD models. Under the constant-rate random-sampling model (case 1), the values are too large to fit in the frame, and they are 737.7 (563.7, 935.7), 629.1 (482.6, 792.4), 494.0 (372.9, 626.4), 482.0 (366.2, 604.2), and 384.3 (291.8, 500.2), respectively. The phylogenetic positions of these nodes are in Supplementary Figure S22.

By relaxing the assumption of constant FBD rates, the estimates of the node ages are quite consistent under diversified and random samplings (Fig. 8, last 5 bars; cf. Table 2). Among them, using 20 time intervals produce slightly younger age estimates than using 10 intervals under both diversified and random samplings. The i.i.d. 20-interval setting has the largest effect on reconciling the ages and reducing DRA; the resulting estimates are very close to those under the constant-rate diversified-sampling FBD model (Fig. 8).

Intuitively, one might wonder if it is enough to divide up time intervals in the middle part of the tree, where most of the fossils are sampled. We thus tested merging the oldest three intervals and the youngest two intervals under i.i.d. rates (i.e., the rate-shifting times are 220, 190, . . ., 70 Ma for *l* = 7). The results are still providing robust age estimates, as we obtained the MRCA age of Hymenoptera as 287.3 (249.8, 329.2) under diversified sampling and 290.7 (254.9, 343.6) under random sampling, although slightly older than the corresponding ages under 10 intervals (see the MCC trees in Supplementary Material). Zhang et al. (2016) employed the SFBD prior with only three time intervals (*d, r, s* shifted at 252 and 66 Ma), but had DRA under random sampling. It turns out that only when sufficient flexibility is allowed in the SFBD process, then the age estimates under different sampling assumptions are reconciled.

It appears that the *λ, µ, ψ* parameterization works better than the *d, r, s* parameterization for reconciling the ages (Fig. 8, 3rd and 4th bars). This behavior may be explained by the parameter *d_i_* being restricted in our analyses to be positive (i.e., *λ_i_ > µ_i_*) across all intervals, but the *λ, µ, ψ* parameterization does not have this restriction, so that *µ_i_* can be larger than *λ_i_*. This gives the latter variant of the model more flexibility in accommodating variation in the speciation, extinction and sampling processes over time. We note that we indeed infer higher extinction than speciation rates under the latter parameterization for some intervals, for example, between 160 and 130 Ma (Figs 9 and 10). However, this pattern is not supported by previous studies of insect diversification dynamics (e.g., Condamine et al., 2016), but seems to be an artifact of fossil-sampling bias. More than half of the hymenopteran fossils are in the Middle and Late Jurassic (about 170–150 Ma), but no fossil is from 150–120 Ma (Supplementary Table S1). This sampling bias somehow is not reflected in the fossil-sampling rate but explained by higher extinction rate.

**Figure 9:**
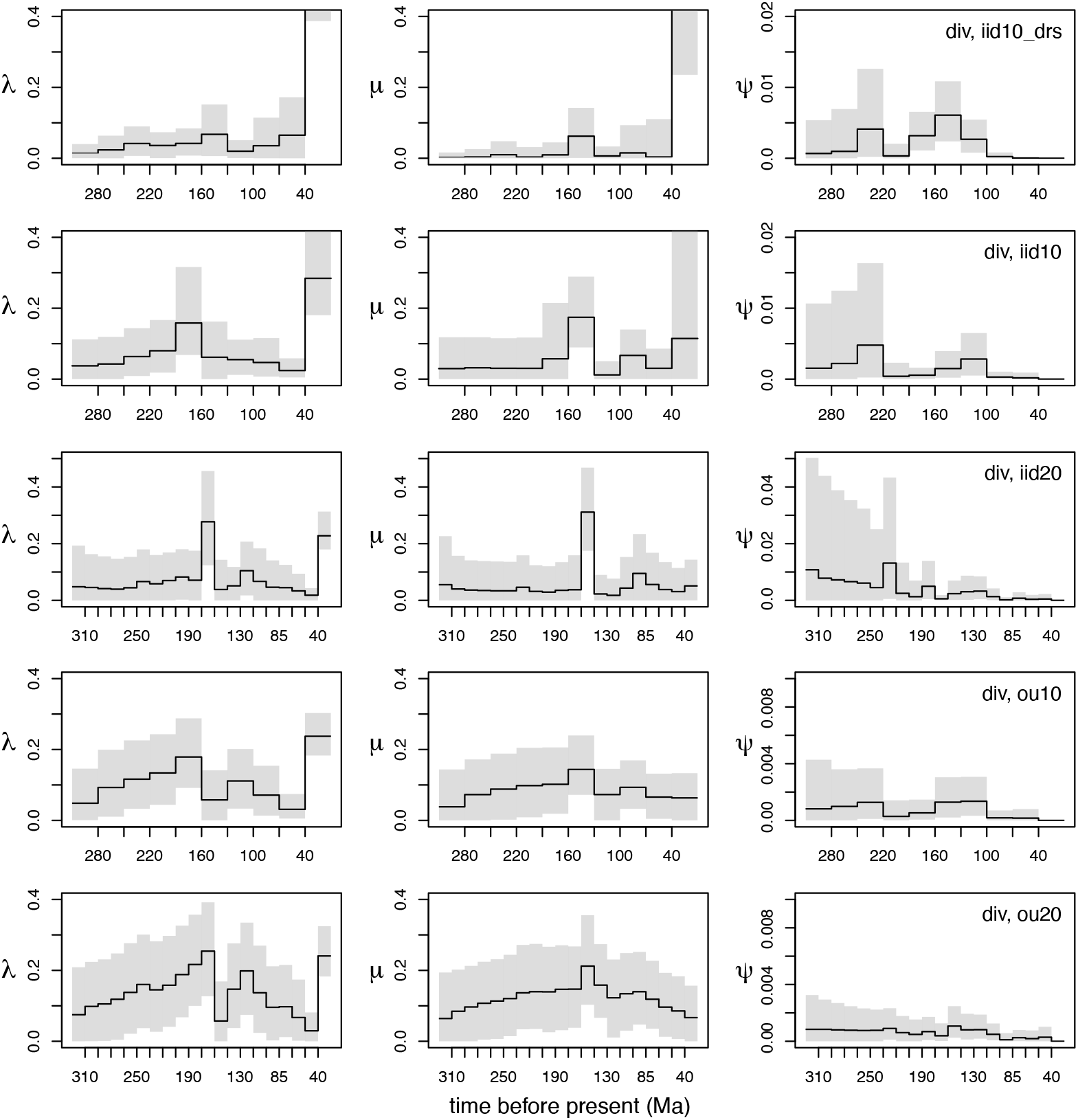
Posterior estimates of the SFBD rates (median and 95% HPD CI) under diversified sampling in the Hymenoptera analyses, using the SFBD model settings shown in Table 2 (Inferences 3–7). The values out of the frame in the last interval in the first two panels are 2.49 (0.39, 6.89) for *λ*_10_ and 2.39 (0.24, 6.84) for *µ*_10_. See Supplementary Figure S23 for the corresponding rates sampled without morphological and molecular data.

**Figure 10:**
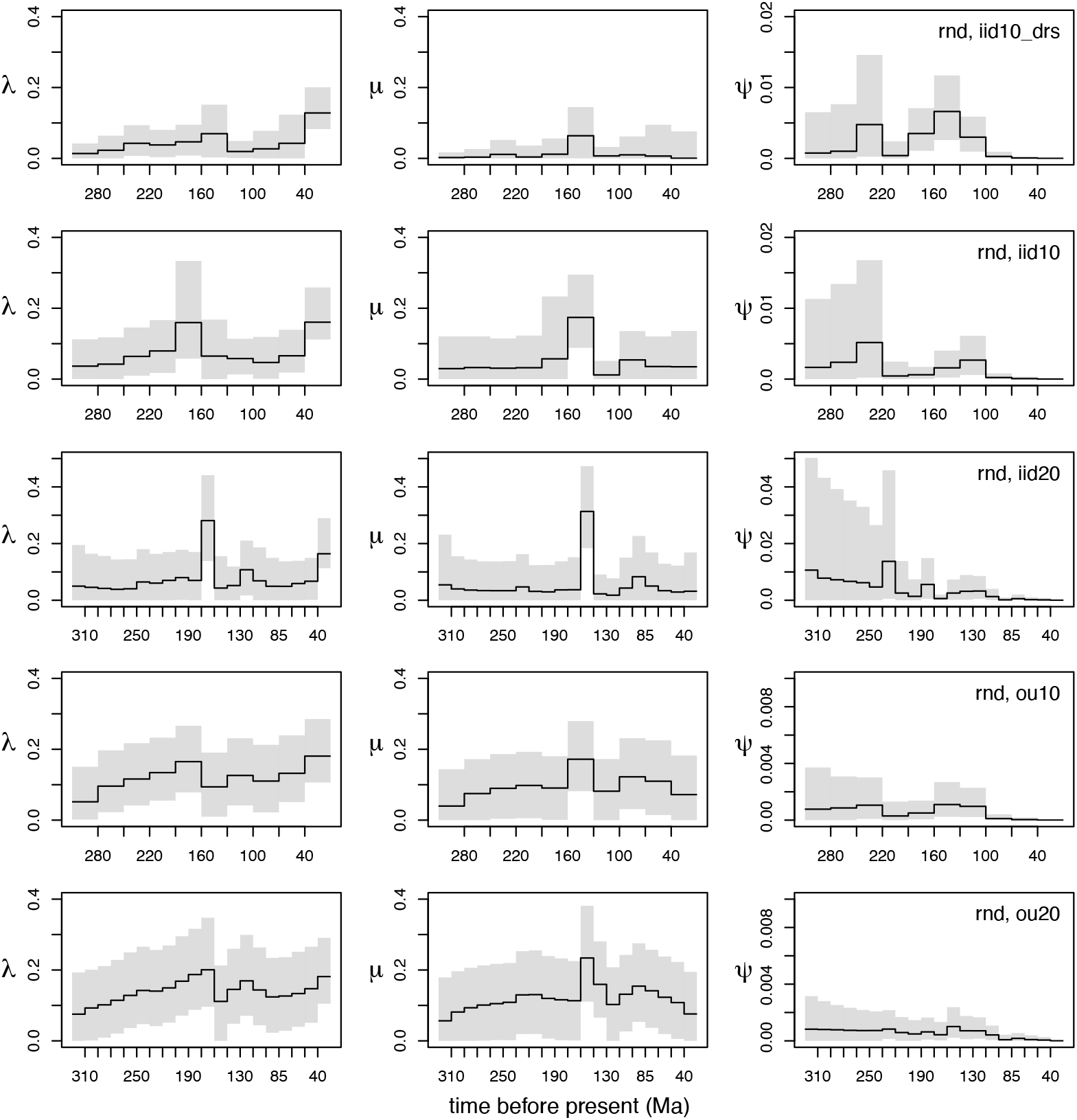
Posterior estimates of the SFBD rates (median and 95% HPD CI) under random sampling in the Hymenoptera analyses, using the SFBD model settings shown in Table 2 (Inferences 3–7). See Supplementary Figure S24 for the corresponding rates sampled without morphological and molecular data.

Taking a closer look at the FBD rates, we find that the rates in the early time intervals are barely affected by the extant-sampling assumptions. They are quite consistent between random and diversified samplings under each prior setting (Fig. 9 vs. Fig. 10, the left 2/3 of each panel). However, the extant sampling assumption can shape the diversification rates quite differently in the most recent intervals (Fig. 9 vs. Fig. 10). This pattern has also been revealed in the simulations under mismatched sampling assumptions (Figs 3 and 5). Under diversified sampling, we see a dramatic rise in speciation rates towards the recent (Fig. 9; note that the *λ* and *µ* estimates for the most recent interval are off the chart under the *d, r, s* parameterization), coupled with a dip a few intervals back in time (except for the *d, r, s* parameterization). The fall and rise of speciation rates towards the recent are less dramatic under random sampling (Fig. 10). The extinction rates are less affected, and the fossil-sampling rates are consistent between diversified and random extant sampling.

The trends are in agreement in general under the i.i.d. and OU priors. But due to the OU smoothing effect, the rates tend to change more gradually over time, and the HPD CIs of the fossil-sampling rates are narrower. This phenomenon is most obvious when time was divided into 20 intervals (Figs 9 and 10). Increasing the hyperprior mean for *σ* (using Gamma(4, 0.2) instead of Gamma(4, 0.05)) produced similar rate variation through time with wider CIs (Supplementary Fig. S25).

### 3.3 Eutheria

With the help of the Boreoeutheria node calibration, we achieved better MCMC convergence and more consistent age estimates under random and diversified sampling, compared with the previous study (e.g., the age of Placentalia was 118 Ma under diversified sampling and *>*300 Ma under random sampling) by Ronquist et al. (2016). Yet still, DRA is apparent under the random-sampling constant-rate FBD model (Fig. 11b, first 2 bars). The SFBD priors resolve DRA under random sampling, resulting in age estimates consistent with diversified sampling (Fig. 11, last 5 bars). As in the Hymenoptera study, the i.i.d. 20-interval setting produces the youngest ages that are close to those under diversified-sampling constant-rate FBD model. Similar to the explanation for the Hymenoptera analyses, different sampling assumptions (diversified vs. random) on extant taxa result in consistent rate estimates in early time intervals but different speciation rates in the most recent time intervals (Figs 12 and 13). Under diversified sampling, the high speciation rate is coupled with high extinction rate in the youngest time interval; while under random sampling, both the speciation and extinction rates are relatively lower in the youngest time interval. Similarly, the OU process priors show the smoothing effect, especially when time was divided into 20 intervals (Figs 12 and 13, Supplementary Figure S29). Given that the diversification rate estimation is sensitive to taxa sampling and phylogeny, we remain cautious interpreting the results.

**Figure 11:**
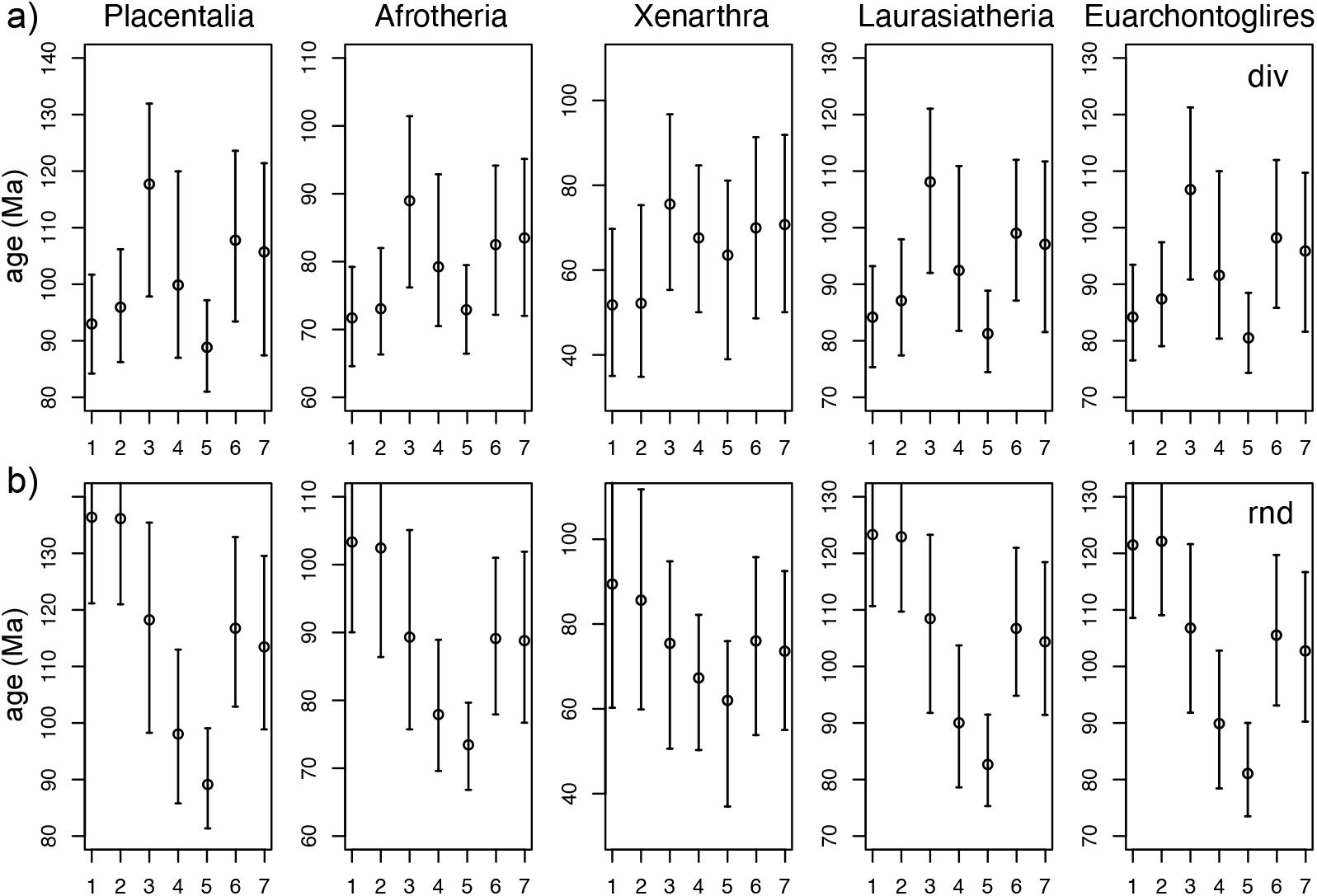
Posterior estimates of five representative node ages (Ma, median and 95% HPD CI) in the Eutheria analyses, using the SFBD model settings shown in Table 2 under a) diversified sampling and b) random sampling. The first two cases in each panel are under constant-rate FBD models. The phylogenetic positions of these nodes are in Supplementary Figure S26.

**Figure 12:**
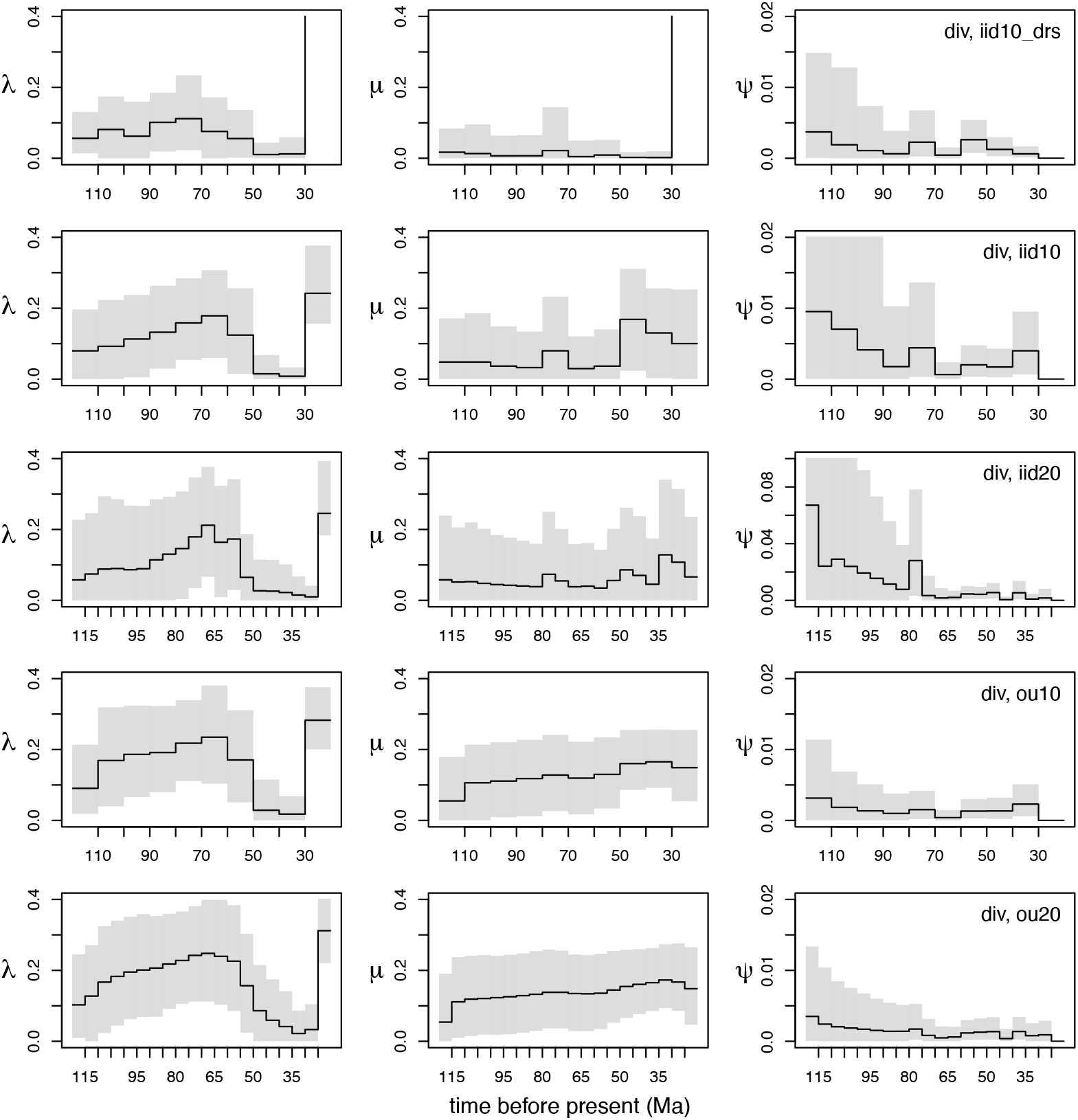
Posterior estimates of the SFBD rates (median and 95% HPD CI) under diversified sampling in the Eutheria analyses, using the SFBD model settings shown in Table 2 (Inferences 3–7). The values out of the frame in the last interval in the first two panels are 1.45 (0.72, 2.44) for *λ*_10_ and 1.39 (0.61, 2.39) for *µ*_10_. See Supplementary Figure S27 for the corresponding rates sampled without morphological and molecular data.

**Figure 13:**
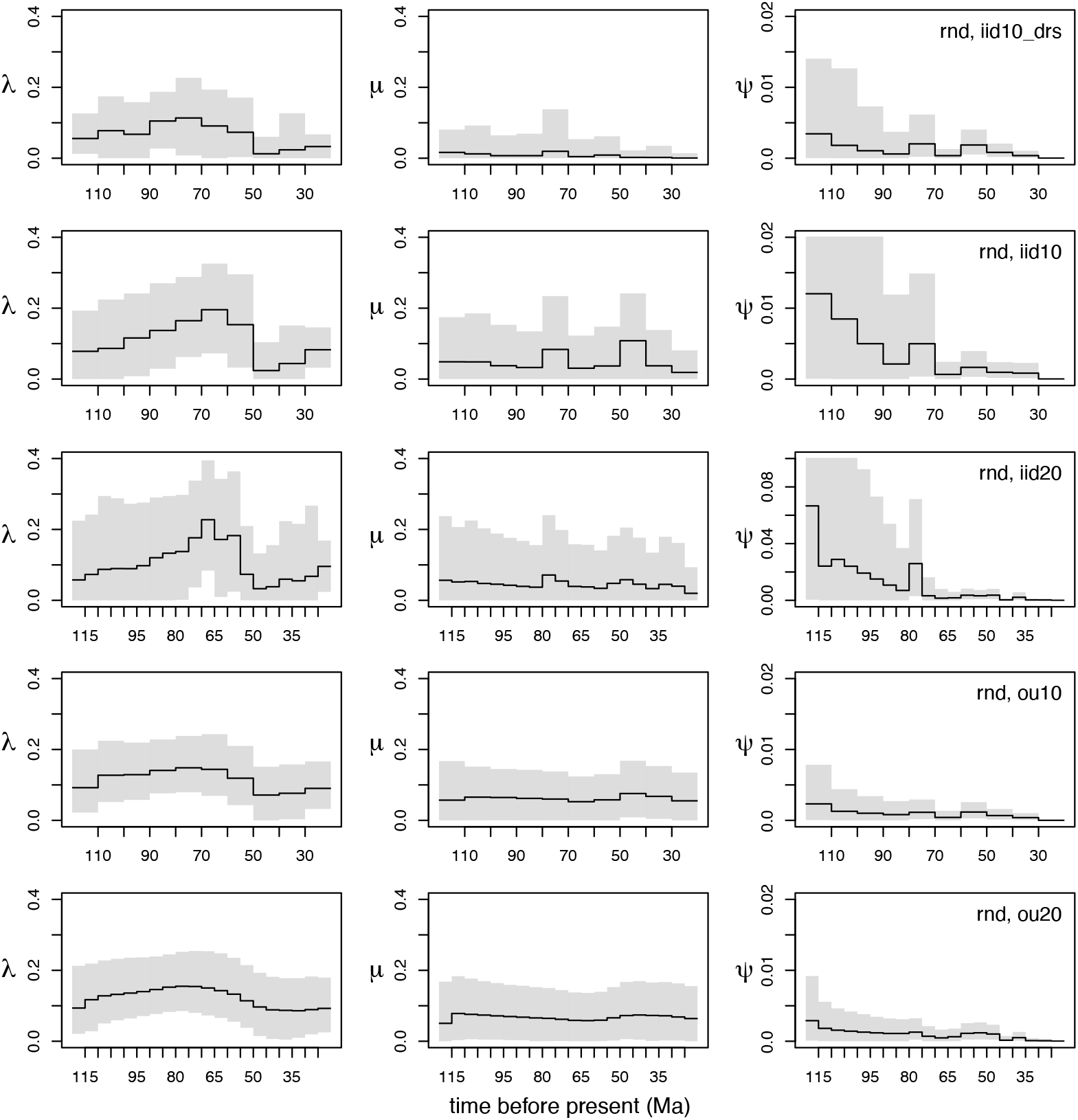
Posterior estimates of the SFBD rates (median and 95% HPD CI) under random sampling in the Eutheria analyses, using the SFBD model settings shown in Table 2 (Inferences 3–7). See Supplementary Figure S28 for the corresponding rates sampled without morphological and molecular data.

## 4 Discussion

The SFBD model used in total-evidence dating is biologically realistic and flexible through explicitly modeling the diversification and sampling processes while allowing rate variations through time and alternative extant sampling assumptions. Our simulations reveal that the root age estimation is robust to various sampling biases, which is achieved by compromising the rate estimates in recent time intervals (e.g., Figs 3 and 5). These patterns shed light on how to interpret empirical results from Hymenoptera and Eutheria analyses, especially the striking differences in birth and death rates toward the present. However, we note that the simulations are designed primarily for validating the implementation of the SFBD model; some simplifications are inevitably involved, specifically, the morphological evolutionary model (Mkv) was perfectly matched between simulation and inference, and constant FBD rates were assumed in simulating the data (although the skyline model with 10 intervals was used for inference). Investigations of the complexities of morphological evolution (such as correlation, irreversibility, convergence, etc), and of variation in speciation, extinction and fossil-sampling rates over time, remain a future task.

For both the Hymenoptera and Eutheria data, these higher-level taxa are arguably more diversely sampled than random sampling, but the diversified-sampling FBD model still is not a perfect match. Take the Hymenoptera data for example, the extant sampling proportion (*ρ* = 0.001) is very small, meaning 99.9% extant divergences (unsampled) must have happened later than 40 Ma (the youngest rate shift), or about 80 Ma at most (the youngest fossil *Prosyntexis*, if we push the rate-shifting time towards the past), which appears highly unlikely. This model mismatch results in an apparent dip and sudden burst in speciation rate towards the present (Fig. 9). If we increase the diversified sampling proportion to 0.1 (Table 2, 4b & 6b), making the sampling model less extreme, the resulting rate estimates are similar to those under random sampling of 0.001, while the ancient age estimates are nearly unchanged (Supplementary Fig. S25). Similar results are also obtained for the Eutheria data under diversified sampling proportion of 0.2 (Supplementary Fig. S29). In summary, interpreting the diversification rate variation in recent intervals should be done with caution, as they are sensitive to the extant sampling assumptions.

The current diversified-sampling method disallows fossil sampling more recent than the cutoff time (*t_c_*). Further, the current implementation also disallows diversification rate shifting after *t_c_* in consideration of lacking data to infer those variable rates, although the mathematical model has been extended to allow such rate variation (Stadler and Smrckova, 2016). The random sampling assumption does not have these restrictions, and theoretically, fossil or rate shift at any time can be included in the SFBD model. However, it does not mean random sampling presents a closer match to the data sampling scheme. Further efforts are needed to develop models that lie in between diversified and random sampling, and relax the fossil sampling assumption, to better capture the biological sampling process.

For phylogenies without fossils (ultrametric trees), we cannot co-estimate *λ* and *µ* due to parameter nonidentifiability (even when *ρ* is fixed as in all of our analyses), if both rates are allowed to vary arbitrarily (Louca and Pennell, 2020). Similar unidentifiable issues exist in serially-sampled trees (Louca et al., 2021). However, Legried and Terhorst (2022, 2023) showed that piecewise-constant birth-death models as used in this paper (albeit also including fossils which is not considered in the theoretical proofs) are identifiable. Nevertheless, even with asymptotic identifiability, Morlon et al. (2022) advise that practical identifiability issues may be common in typically sized phylogenies. Thus, carefully determined constraints or regulations (e.g. through informative or smoothing priors as done in this study) are typically required.

A smoothing prior can be used to model correlations among neighbor-hood intervals, thus relieving the practical unidentifiability that would arise when using i.i.d. rates and many time intervals, as an individual interval would have very little information to infer independent rates. Magee et al. (2020) was able to utilize more than a hundred intervals through properly regulating the Horseshoe Markov random field prior. In contrast to Brownian motion, the OU process has both diffusion and mean-reverting (drift towards its mean) effect. One complication of the OU prior (and other smoothing priors in general) is that it is parameter rich, and specifying hyperpriors for these hyperparameters can be tricky. A general principle is to use diffuse distributions to avoid enforcing strong trends. The distributions we used in this study appear to be flexible enough (Supplementary Fig. S1), which is an indication that they could be a good default choice. On the other hand, if the user has strong belief in a rate (e.g., speciation rate) to be increasing or decreasing through time (e.g., because there is an increase or decrease in net species diversity in the fossil record over time), it is possible to adjust the distributions for the initial state *x*_0_ and long-term mean *θ* to be more informative so that they reflect such an expectation. In practice, with moderate size of data (hundreds of tips or less), it is likely that the posterior estimates are sensitive to the prior. It is important to be aware of this when using informative priors. It is always good practice in Bayesian analyses to explore the effect of different prior assumptions on the posterior. In some cases, it might be appropriate to use more than 20 intervals under the OU model, or even relaxing its univariate assumption by allowing *ν* and/or *σ* to vary over time, but the effect of regularizing the hyperparameters and the statistical power in detecting shifts in their values require further exploration.

At the current stage, the SFBD implementation requires predefining the number and shifting time of the intervals. For the empirical data sets and model settings examined here, using 10 to 20 uniformly spaced intervals works pretty well. Generally, the appropriate number of intervals should depend on the size of the tree, with more intervals feasible for larger trees. Ultimately, it would be tempting to automatically select the number and shifting time of the intervals, as well as to jump between diversified and random sampling assumptions, using a model averaging technique such as reversible-jump MCMC (Green, 1995). However, making this work would require a significant amount of technical redesign, programming and testing, which is beyond the scope of this study. An alternative would be to use model selection based on marginal likelihoods and Bayes factor (Kass and Raftery, 1995). However, this is computationally very demanding and does not seem feasible for these data sets with the currently available techniques. Importantly, such model comparison is inevitably sensitive to the prior settings, so the results need to be carefully evaluated with this in mind.

In this study, we have revealed that total-evidence dating under the SFBD model with both i.i.d. and OU priors is robust to extant sampling assumptions. For the two total-evidence data sets, allowing flexible variation over time in speciation, extinction and fossil-sampling rates in the SFBD model pushed the dramatically old age estimates observed under the constant-rate FBD model with random sampling towards the present, and greatly reduced the gap between the root age and the age of the oldest fossil of the focal taxa. In particular, the tree ages are similar for very different extant-species sampling assumptions.

The robustness of the SFBD model is due to its flexibility, allowing it to absorb misfit in the fossil and extant sampling assumptions by adjusting the speciation, extinction and fossilization rates of particular time intervals. As our sampling assumptions improve, we expect the SFBD model to become increasingly informative about true macroevolutionary dynamics. The SFBD model with more realistic sampling assumptions will be instrumental in inferring macroevolutionary processes accurately from total-evidence data sets.

## 5 Software Availability

The SFBD model with diversified sampling and OU smoothing prior has been implemented in a modified version of BDSKY available from https://github.com/zhangchicool/bdsky, which supports BEAST version 2.6. It has been merged to the main branch of BDSKY and will support BEAST version 2.7 and higher in the near future. At the moment, many SFBD settings cannot be set up in BEAUTi. To use diversified extant sampling, the user has to modify the following block in the XML file generated from BEAUTi, from

`spec="beast.evolution.speciation.BirthDeathSkylineModel"` to

`spec="beast.evolution.speciation.BirthDeathSkylineDiversifiedSampling"`.

In Supplementary Material, we further provide xml files showing how to set the OU prior for the SFBD rates. The shifting times are specified independently for each rate, allowing each rate to shift at different time points.

Here we also mention the relevant commands in MrBayes, which are applicable to the developmental version at https://github.com/NBISweden/MrBayes. These features will be included in version 3.2.8 in the near future. Different from version 3.2.7 and described in Zhang et al. (2016), the shifting times are specified independently for each rate, in the order of fossil-sampling, net diversification and turnover. For example, this command specifies the 10 epochs in the Hymenoptera analyses: `"prset samplestrat = diversity 9: 280 250 220 190 160 130 100 70 40, 9: 280 250 220 190 160 130 100 70 40, 9: 280 250 220 190 160 130 100 70 40;"`, whereas using `"prset samplestrat = diversity 2: 252 66;"` allows the fossil-sampling rate to shift twice while keeping the speciation and extinction rates constant. A NEXUS file containing all the MrBayes commands for doing the comparison with BEAST 2 can be found in Supplementary Material.

## 6 Supplementary Material

Supplementary files can be found in data repositories, Zenodo (https://doi.org/10.5281/zenodo.8145628) and Dryad (https://doi.org/10.5061/dryad.573n5tb41).

## Acknowledgments

We thank Louis du Plessis for his generous help on the BDSKY implementation and on using the R-package bdskytools, and thank Sebastian Höhna and four anonymous reviewers for their constructive comments. This research was funded by the National Natural Science Foundation of China (42172006) and the Hundred Young Talents Program of Chinese Academy of Sciences (Y902061), both to C.Z. Early work of this study was also supported by the European Research Council under the Seventh Framework Programme of the European Commission (PhyPD: grant number 335529 to T.S.). F.R. acknowledges the Swedish Research Council (grant 2018-04620). Computations were performed using the cluster in the High Performance Computing Center of Institute of Vertebrate Paleontology and Paleoanthropology.

